# Autophagy Dysfunction in iPSCs-Derived Neurons and Midbrain Organoids Carrying a *SNCA* Triplication

**DOI:** 10.1101/2025.08.08.669314

**Authors:** Catarina Serra-Almeida, Javier Jarazo, Gemma Gomez-Giro, Isabel Rosety, Alise Zagare, Daniele Ferrante, Cláudia Saraiva, Daniela Frangenberg, Elisa Zuccoli, Ana Clara Cristóvão, Liliana Bernardino, Jens Christian Schwamborn

## Abstract

Parkinson’s disease (PD), characterized by α-Synuclein aggregation and dopaminergic neuronal loss, has no current cure. Autophagy is critical for α-Synuclein clearance, yet its real-time dynamics remain challenging to assess in human-relevant systems. Here, we used live-cell imaging to assess autophagy within human neuronal cultures and midbrain organoids (hMOs) derived from induced pluripotent stem cells (iPSCs) of PD patients carrying a triplication of the α-Synuclein gene (3xSNCA). Using the LC3-Rosella dual-fluorescent reporter, we quantified autolysosomes dynamics in real time. In 3xSNCA neuronal cultures, we detected early autophagy defects. In 3xSNCA hMOs, reduced autolysosome area, increased total and phosphorylated α-Synuclein (pS129), and decreased electrophysiological activity were observed at 50 days of differentiation (DoD). By 70 DoD, autophagy impairment became more pronounced, overlapping with dopaminergic neuron loss. These findings support the use of human iPSCs-derived models to study autophagy dysfunction in PD and demonstrate a temporal correlation between impaired autophagy, α-Synuclein pathology and neuronal degeneration.

## Introduction

Affecting nearly 10 million people globally, Parkinsońs disease (PD) is a neurodegenerative condition that progresses over time. It belongs to the synucleinopathies, a group of disorders marked by abnormal accumulation of α-Synuclein in several brain areas. Encoded by the SNCA gene, this small protein is mostly found in the presynaptic terminals of neurons, where it plays a role in neurotransmitter release ^1,2^.

*SNCA* point mutations, such as A53T, A30P, and E46K, as well as gene duplications and triplications are linked with familial PD forms ^3–8^. In particular, *SNCA* triplication (3xSNCA) is correlated with elevated α-Synuclein levels and early-onset forms of PD ^7,9,10^. Even in idiopathic PD in which genetic mutations have not been identified, α-Synuclein accumulation is observed. From this perspective, studying its regulation and clearance mechanisms is crucial to better understand disease mechanisms and identify new treatment avenues ^11^.

Autophagy is a key cellular process in the regulation of α-Synuclein levels. It includes a number of pathways such as chaperone-mediated autophagy (CMA) and macroautophagy, both of which are implicated in PD pathogenesis ^12,13^. CMA depends on specific proteins like heat shock cognate 70 (HSC70) and lysosomal-associated membrane protein 2A (Lamp2A) to degrade cytosolic proteins containing the KFREQ motif including wild-type α-Synuclein ^14^. Yet, mutant α-Synuclein impairs this pathway which leads to macroautophagy as a compensatory mechanism ^15,16^. Macroautophagy involves the delivery of cellular waste, including misfolded or aggregated proteins like α-Synuclein, to lysosomes, where acidic enzymes are responsible for their breakdown. This process depends on the fusion of autophagosomes (double-membraned vesicles) with lysosomes to form autolysosomes ^12^.

The accretion of α-Synuclein aggregates together with other pathological mechanisms, such as lysosomal dysfunction, contribute to the degeneration of dopaminergic neurons in *substantia nigra pars compacta*, which are particularly vulnerable ^17^. This interaction is interesting since it seems to show a two-way link. The accumulation of α-Synuclein can hinder autophagy, hence facilitating more its aggregation.

Evidence from post-mortem PD brains points to defective autophagy, showing increased autophagosomes, abnormal lysosomes, and high LC3II levels. Lewy bodies positive for LC3 have also been detected in the *substantia nigra* ^18^. Furthermore, reduced levels of HSC70 and Lamp2A have also been observed ^19^. Similar findings were observed in animal models ^20^. Studies have shown that lysosomal defects can impair macroautophagy, leading to the buildup of α-Synuclein ^21,22^. These results confirm that autophagy is disrupted in PD, across models and species. Still, little is known whether these impairments are a cause or a consequence of α-Synuclein accumulation.

Therefore, it is important to unravel this relationship by identifying human-relevant models that allow real-time tracking of autophagic dynamics. Advances in induced pluripotent stem cells (iPSCs) technology led to the development of patient-specific neuronal cell culture and organoid models that replicate key features of PD ^23^. Although iPSCs-derived neuronal cultures provide a practical and accessible model for PD modelling ^24^, their two-dimensional (2D) architecture lacks cell-cell and cell-extracellular matrix (ECM) interactions and midbrain complexity ^25,26^. In contrast, three-dimensional (3D) organoids are a more advanced system that can self-organize into structures that resemble the architecture and function of human organs ^27^. Specifically, human midbrain organoids (hMOs) can generate a large number of dopaminergic neurons and glial cells, and release dopamine ^28,29^. Several studies have used iPSCs-based cell models to explore *SNCA*-related mechanisms in PD ^30^. Our research group, alongside others, has shown that iPSCs-derived hMOs carrying a 3xSNCA show dopaminergic neuron loss and α-Synuclein aggregation ^31–34^. This evidence indicates their relevance as PD models.

However, even with growing evidence of autophagy disruption in PD, most studies rely on static endpoints analysis, which do not capture the progression and the timing of the impairments. Hence, the temporal dynamics of autophagy in human 3xSNCA models remain poorly understood, representing a major gap for the field. To tackle this limitation, we implemented the LC3-Rosella system, a dual-fluorescent construct enabling the visualization of autolysosomes in live cells. By applying this tool to both neuronal cultures and hMOs derived from iPSCs, we aimed to investigate how autophagy changes throughout disease progression in these models.

In neuronal cultures, autophagy exhibits dynamic modulation over the course of time. Autophagic activity was found decreased in the beginning of neuronal differentiation, followed by transient increase and subsequent decline. In contrast, PD hMOs exhibited a progressive and sustained reduction in autophagy, particularly at 70 days of differentiation (hereafter referred to as DoD). PD hMOs also exhibited increased total and phosphorylated α-Synuclein (pS129) in addition to its aggregation at these later timepoints (50 and 70 DoD), followed by dopaminergic neuron loss. This temporal correlation suggests that autophagy failure may precedes and possibly contributes to pathological α-Synuclein inclusions. Altogether, these findings reinforce the link between autophagy impairment and α-Synuclein pathology.

## Results

### PD neuronal cultures show early and progressive autophagy impairment

This study is based on several human iPSCs lines, as detailed in Supplementary Table 1. The PD patient line, carrying a 3xSNCA, was edited to incorporate the autophagy sensor (PD-auto), created by integrating the LC3-Rosella sensor ^35^. For comparison, we included two sex- and age-matched healthy control lines: a standard control (Control) and a modified control (Control-auto) that also incorporates the autophagy sensor. We previously showed that all iPSCs lines exhibited embryonic stem cell-like morphology and expressed pluripotency markers (SOX2, OCT4, NANOG) ^33^.

iPSCs were differentiated into neuroepithelial stem cells (NESCs) with a hindbrain/midbrain identity, as we previously described ^36^. To confirm neural stem cell identity, NESCs were characterized by the expression of neural progenitor markers, including SOX1, SOX2, Nestin and PAX6, which indicates dorsal identity (Supplementary Fig. 1; see Supplementary Table 2 for the list of antibodies). Neuronal cultures were then generated from these NESCs and maintained for up to 11 DoD. The schematic representation of neuronal cultures generation is in Supplementary Fig. 2.

To investigate early autophagy dynamics, we first analyzed iPSCs-derived 2D neuronal cultures using the LC3-Rosella autophagy sensor, which incorporates the DsRed and pHluorin fluorescent proteins. In acidic environments, like the autolysosome lumen, pHluorin is quenched while DsRed remains fluorescent, indicating fusion with lysosomes ^35^. This allowed us to identify autolysosomes that appeared as red vesicles (Fig. 1a). Representative images in Fig. 1b1 and 1b2 show the progressive reduction in red-only vesicles in PD neurons, indicative of impaired autophagy over time. We assessed the area and density of autolysosomes, which were further classified as small, medium, and large, based on their size distribution. The size of these structures, described in the methods section, can be used as an indicator of their functional state. Small autolysosomes are usually more efficient in degrading materials, while larger ones suggest decreased degradation capacity ^37,38^.

**Figure 1.**
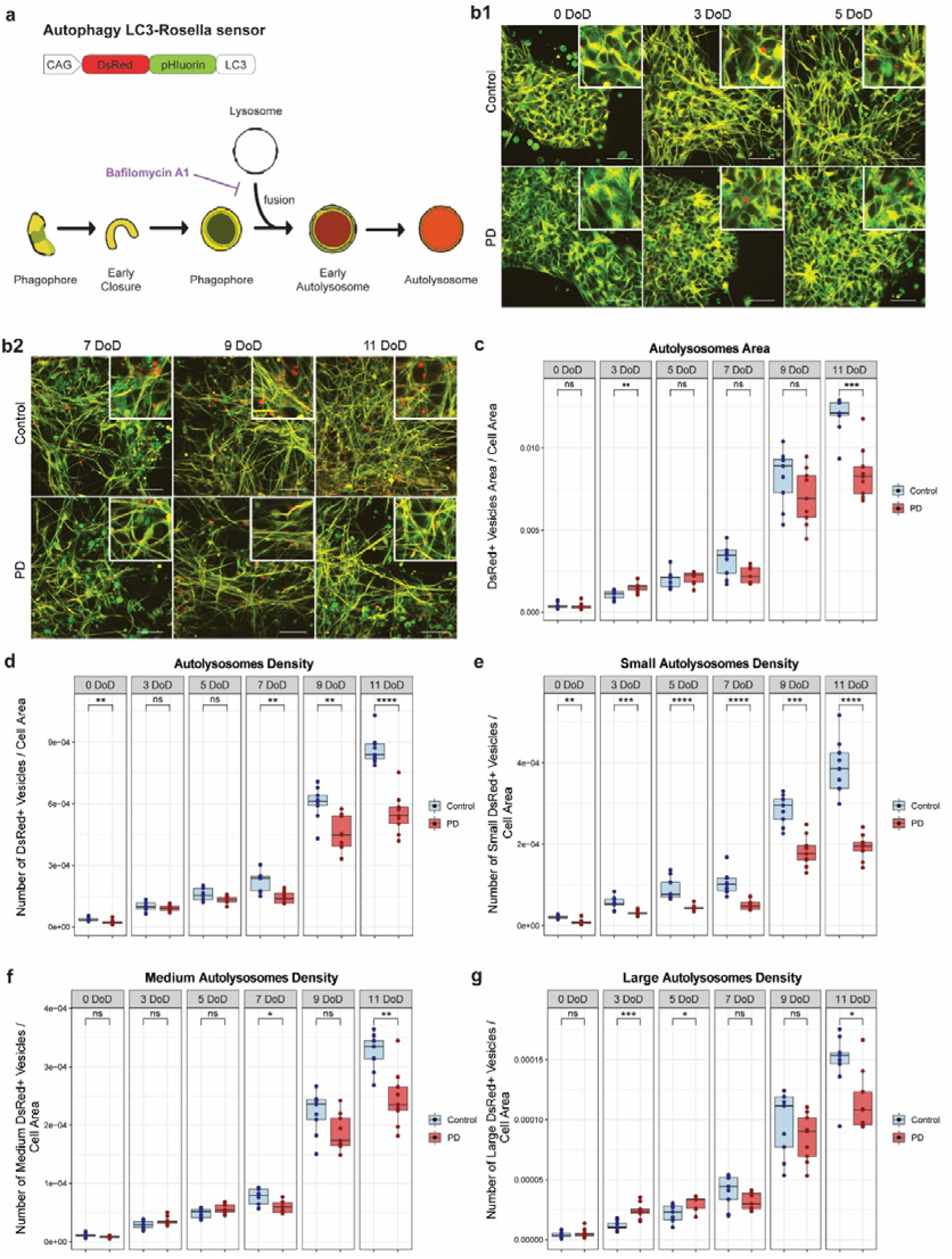
Autolysosome size and density in PD neurons over 11 DoD. (a) Structure of LC3-Rosella sensor, using the pH-sensitive pHluorin and DsRed proteins. It allows us to distinguish autolysosomes (pHluorin-DsRed+ structures) in our cellular model. (b1, b2) Representative live-cell microscopy images of neuronal cultures expressing the LC3-rosella autophagy reporter. Images show control and PD neurons at all stages of differentiation, with their respective zoomed region of interest. Scale bar: 20 μm. (c-g) Quantitative analysis of autophagy in neuronal cultures using the LC3-rosella reporter, at 0, 3, 5, 7, 9 and 11 DoD. PD neurons showed dynamic autophagy impairment over time compared to control. (c) Total autolysosome area per cell area. (d) Number of autolysosomes per cell area. (e-g) Number of small (e), medium (f), and large (g) autolysosomes per cell area. Data result from three independent experiments each analyzed in triplicate. Values are normalized to the average of controls across all time points, allowing a comparison over time. Statistical analysis was performed using Wilcoxon T-test; *p < 0.05, **p < 0.01, ***p < 0.001, and ****p < 0.0001, with “ns” indicating non-significant results. Image analysis was performed using a custom MATLAB script for automated quantification of the different parameters of the autolysosome analysis.

At 0 DoD, total autolysosome area was similar, but PD cultures had fewer small autolysosomes (Fig. 1c-e), suggesting early degradation defects. No differences were observed in the density of medium and large autolysosomes at this stage (Fig. 1f, g). At 3 DoD, PD cultures exhibited a higher autolysosome area compared to controls, with no difference in the overall autolysosome density (Fig. 1c, 1d). At the same time, PD neurons had more large autolysosomes (Fig. 1g) but fewer small ones (Fig. 1e), that might indicate an accumulation of autolysosomes that failed to degrade their contents properly. At 5 DoD, no differences were observed in the total autolysosome area and the density of medium autolysosomes in PD neurons (Fig. 1c, 1f). However, PD neurons showed a persistently lower density of small autolysosomes and a higher density of large autolysosomes (Fig. 1e, 1g). By 7 DoD, while the total autolysosome area showed no significant difference between PD and control cultures, we observed a lower autolysosome density across all size categories in PD neurons (Fig. 1c-g), indicating that autophagic activity worsened over time. At late time points of differentiation (9 and 11 DoD), PD cultures showed reduced autolysosome area and density across all sizes in comparison to controls (Fig. 1c-g).

These data indicate progressively impaired autophagy in PD neurons, which begins early and worsens over time. These findings highlight that targeting autophagy, especially at early stages, may prevent PD progression.

### PD hMOs recapitulate autophagy dysfunction observed in PD neuronal cultures

Having established the presence of early autophagic impairment in neuronal cultures, we next sought to assess whether similar alterations occurred in hMOs, a more physiologically relevant context. These 3D structures mimic several aspects of human midbrain and cellular organization, making them suitable for studying long-term autophagic dynamics ^27–29,33^.

For this, we assessed autophagy in mosaic hMOs at 50 and 70 DoD by live imaging. We focused on 50 and 70 DoD, as they represent intermediate and late stages of maturation, respectively. The schematic representation of hMOs generation is detailed in Supplementary Fig. 2. Representative images of control and PD mosaic hMOs from 70 DoD are shown in Fig. 2a, where PD organoids display a reduction in the autolysosomes.

**Figure 2.**
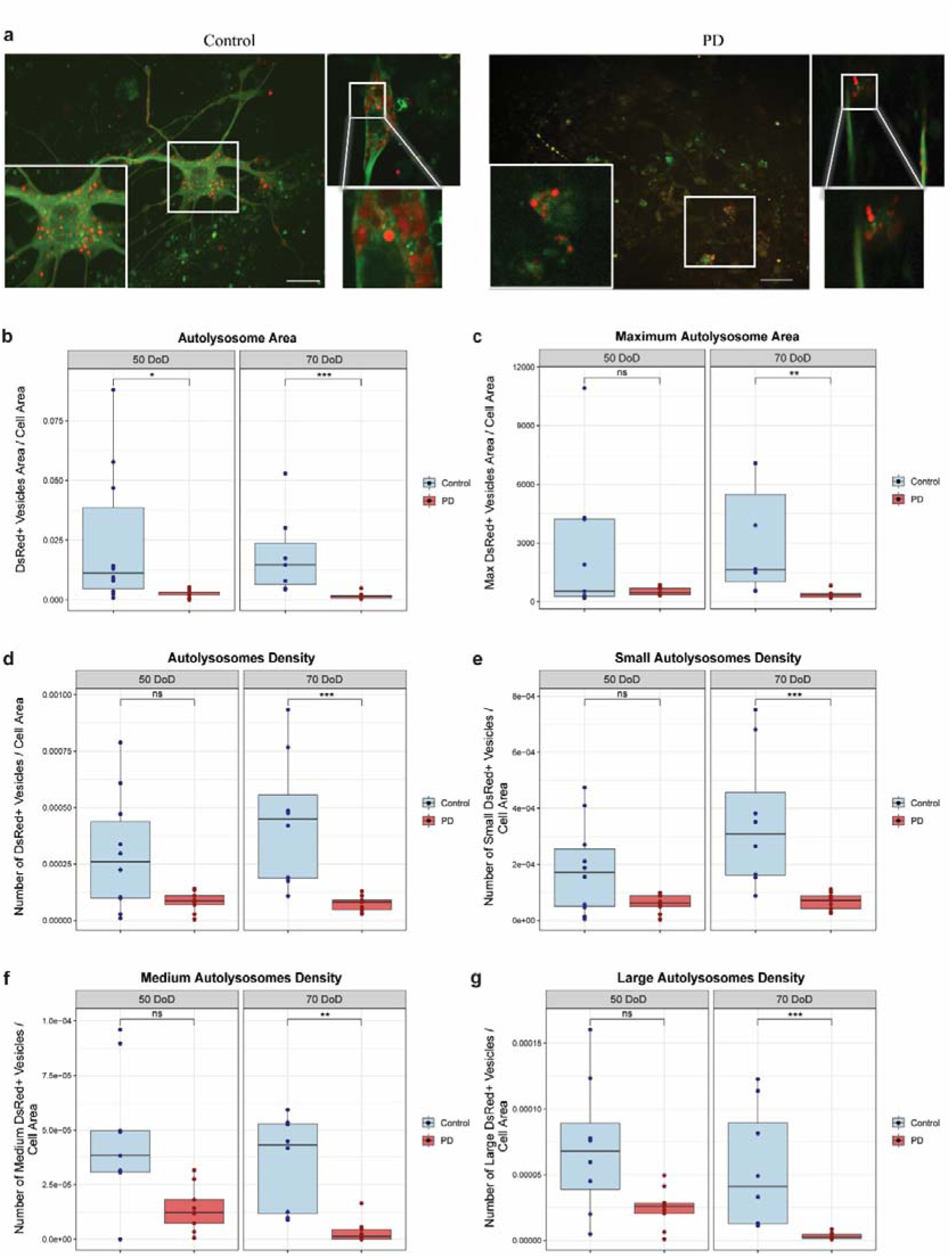
PD hMOs showed autophagy dysfunction. (a) Representative live-cell microscopy images of mosaic hMOs (55% LC3-Rosella expressing NESCs, 45% regular NESCs) at 70 DoD. Images show control and PD mosaic hMOs, with their respective zoomed region of interest. Scale bar: 20 μm. (b-g) Quantitative analysis of autophagy in hMOs using the LC3-Rosella reporter at 50 and 70 DoD. PD mosaic hMOs showed significant autophagy impairment in comparison to controls. (b) Total autolysosome area per cell area. (c) Maximum autolysosome area per cell area. (d) Number of autolysosomes per cell area. (e-g) Number of small (e), medium (f), and large (g) autolysosomes per cell area. Data represent results from three independent experiments, with three fields per hMOs. Values are normalized to the average of controls across both DoD. Statistical analysis was performed using Wilcoxon T-test; *p < 0.05, **p < 0.01, and ***p < 0.001, with “ns” indicating non-significant results. Image analysis was performed using a custom MATLAB script for automated quantification of the different parameters of the autolysosome analysis.

Similar to neuronal cultures, we quantified the area and density of autolysosomes, categorizing them into small, medium, and large sizes based on their areas. Additionally, we analyzed the maximum autolysosome area, representing the largest autolysosome present in each condition. Live imaging of Rosella-LC3 in mosaic hMOs showed that PD hMOs had decreased autolysosome area at 50 and 70 DoD (Fig. 2b). By 70 DoD, all autophagy parameters were reduced. PD hMOs presented reduced maximum autolysosome area, suggesting a possible dysfunction in the fusion or expansion of autolysosomes that are critical for proper cellular degradation (Fig. 2c). They also presented lower autolysosome density across all size categories when compared to controls, indicating a defect in the formation or maturation of autolysosomes at late stages of maturation (Fig. 2d-g).

### PD hMOs exhibit elevated levels of **α**-Synuclein and pS129

Building on our previous findings showing autophagy dysfunction in both PD models, we next investigated whether these impairments correlate with α-Synuclein accumulation and phosphorylation, two hallmark features of PD pathology.

First, we investigated the levels of α-Synuclein, pS129 and the pS129 / α-Synuclein ratio in hMOs at 50 and 70 DoD by Western Blot. Representative immunoblots are presented in Fig. 3a. PD hMOs showed increased levels of total α-Synuclein compared to controls at both time points (Fig. 3b). PD hMOs also had higher levels of pS129 and an increased pS129/α-Synuclein ratio (Fig. 3c, 3d).

**Figure 3.**
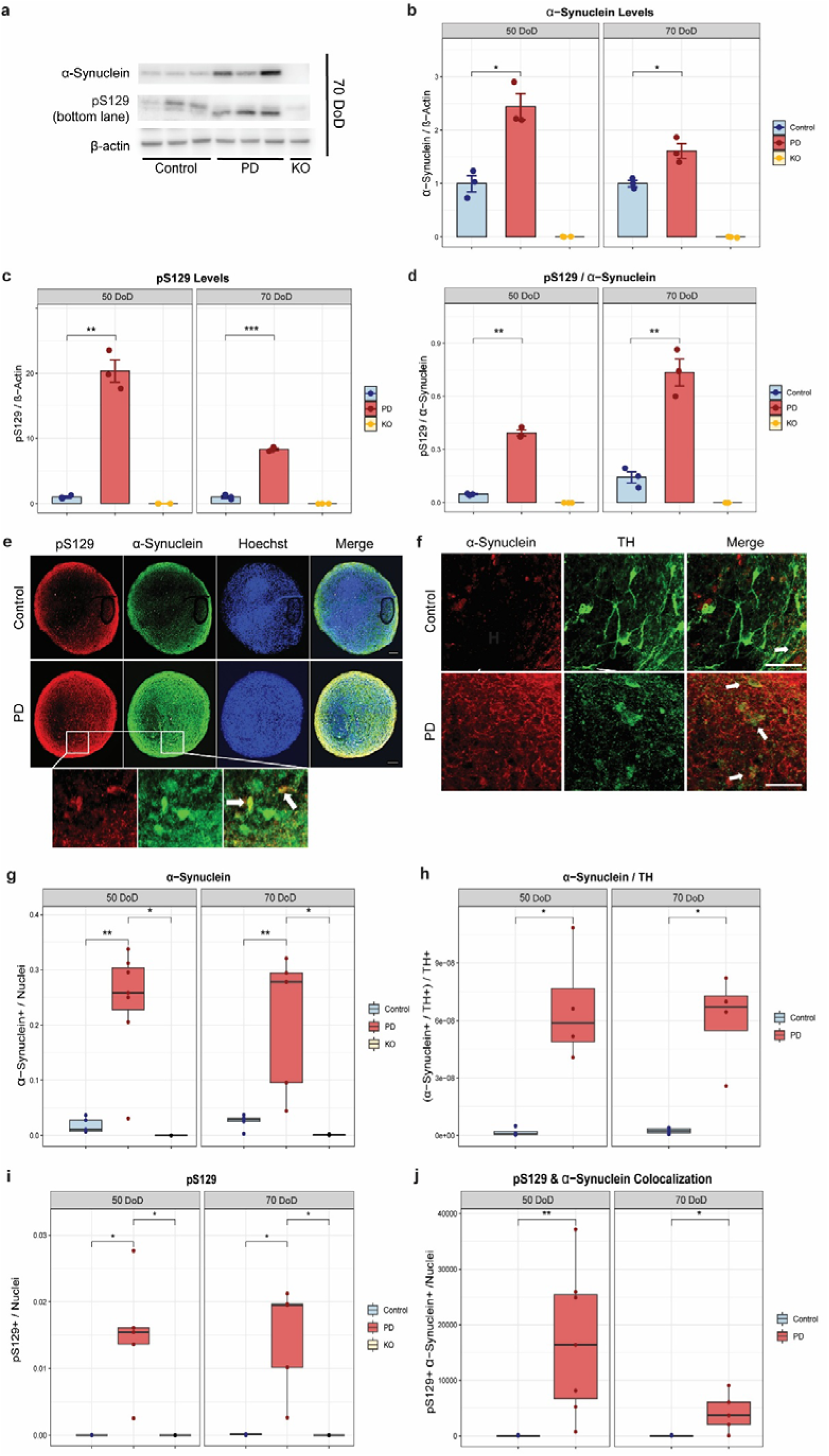
PD hMOs exhibit increased α-Synuclein and pS129 levels. (a) Representative Western blot images showing total α-Synuclein and β-Actin in control, PD and SNCA knockout hMOs, which were used to validated the specificity of the antibodies for α-Synuclein and pS129. (b-d) Bar graphs depict the protein levels of (b) total α-Synuclein, (c) pS129, and (d) the ratio of pS129 to total α-Synuclein where they are increased in PD hMOs. Protein expression was normalized to β-Actin. Error bars represent standard error of the mean (SEM). Immunoblot analysis was performed using ImageJ software. Data represent results from three independent experiments. Values are normalized to the average of controls at each time point. Statistical analysis was performed using Student’s t-test; *p < 0.05, with “ns” indicating non-significant results. (e, f) Representative immunofluorescence images of hMOs stained for pS129, total α-Synuclein, TH and nuclei. Arrows point to double-positive cells. Scale bar: 200 μm. (g, h) Quantification of (g) total α-Synuclein and (h) ratio of total α-Synuclein to TH, revealing accumulation of α-Synuclein in dopaminergic neurons in PD hMOs. (i-j) Quantification of (i) pS129, and (j) colocalization of total and pS129, showing increased phosphorylation of α-Synuclein in PD hMOs. Data result from three independent experiments, each analyzed from at least two fields per hMO. Values are normalized to the average of controls across all time points, allowing a comparison over time. Statistical analysis was performed using Wilcoxon T-test; *p < 0.05, **p < 0.01, with “ns” indicating non-significant results. Image analysis was performed using a custom MATLAB script for automated quantification of the different stainings. The absence of staining in SNCA knockout sections validated the specificity of the antibodies for α-Synuclein and pS129.

To confirm and localize these findings at the cellular level, we performed immunofluorescence stainings. Representative images illustrated in Fig. 3e and 3f, show a widespread of α-Synuclein and pS129 in PD hMOs. The quantification confirmed that PD hMOs had more total α-Synuclein, particularly within TH-positive neurons (Fig. 3g, 3h), implying an accumulation of α-Synuclein in this neuronal population. Consistent with our Western Blot findings, immunofluorescence analysis also showed an increase in pS129 levels in PD hMOs (Fig. 3i). Both the pS129/α-Syn colocalization (Fig. 3j) and the ratio of pS129 to total α-Synuclein (see Supplementary Fig. 3a) were increased in PD hMOs. The elevated levels of both total and pS129 demonstrate that these models recapitulate key pathological features of PD.

### PD hMOs exhibit increased **α**-Synuclein aggregation

Following our observation of elevated α-Synuclein and pS129 levels in PD hMOs and autophagy dysfunction, we investigated whether α-Synuclein aggregates are present in hMOs, at 50 and 70 DoD. Given that α-Synuclein aggregation is a hallmark of PD pathology, we performed immunostainings for markers of protein aggregation, including p62, Amytracker and Proteostat together with α-Synuclein, at 70 DoD (Fig. 4a). Lower magnification images can be found in Supplementary Fig. 3b. The Proteostat® Aggresome Detection Kit is a fluorescent dye that specifically binds to aggregated proteins, while the Amytracker 680 is a red dye that binds to protein aggregates with repetitive arrangement of β-sheets, labeling α-Synuclein aggregates.

**Figure 4.**
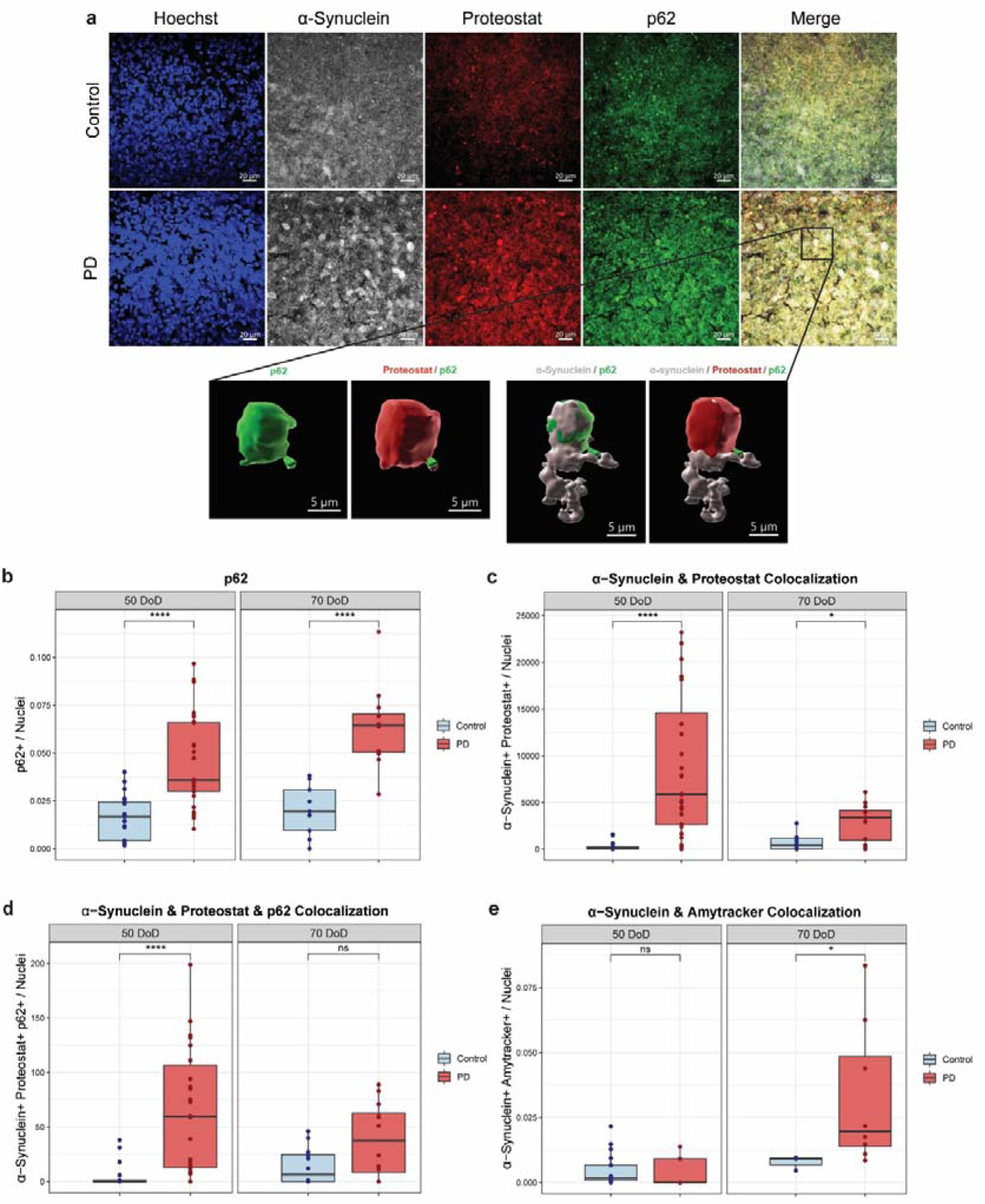
Increased α-Synuclein aggregation in PD hMOs. (a) Representative immunofluorescence images of hMOs stained for α-Synuclein (grey), p62 (green), Proteostat (red) and nuclei (blue) at 70 DoD as well as a 3D reconstruction. Scale bar: 5 and 20 μm. (b-e) Quantification of (b) p62 immunoreactivity, (c) Colocalization of α-Synuclein and Proteostat, (d) triple colocalization between α-Synuclein, Proteostat, and p62 and (e) colocalization of α-Synuclein and AmyTracker in control and PD hMOs, at 50 and 70 DoD, where PD hMOs presented increased levels of aggregation markers. Data represent results from three independent experiments, each analyzed from at least three fields per hMO. Values are normalized to the average of controls across all time points, allowing a comparison over time. Statistical analysis was performed using Wilcoxon T-test; *p < 0.05, **p < 0.01, ***p < 0.001, and ****p < 0.0001, with “ns” indicating non-significant results. Image analysis and colocalization quantification were performed using a custom MATLAB script for automated quantification.

In PD hMOs, p62 levels were elevated compared to controls at both times (Fig. 4b), which might indicate an accumulation of autophagosomes that are not properly fused with lysosomes. Besides being a marker of autophagy, p62 is a common component of protein aggregates in neurodegenerative diseases, including PD ^39^, further confirming impaired clearance of protein aggregates. To specifically detect α-Synuclein aggregates, we performed colocalization analyses using the Proteostat and the Amytracker. We found that PD hMOs showed increased co-localization of α-Synuclein and Proteostat (Fig. 4c) as well as enhanced colocalization of α-Synuclein, Proteostat, and p62 (Fig. 4d), suggesting that while α-Synuclein is being targeted for autophagic clearance, it is not efficiently degraded. Moreover, we found higher colocalization between α-Synuclein and Amytracker in PD hMOs at 70 DoD (Fig. 4e), suggesting the presence of β-sheet-rich α-Synuclein aggregates, that are highly toxic and more resistant to degradation^40^.

### PD hMOs show dopaminergic neuronal degeneration and functional decline

Lastly, we investigated whether hMOs undergo dopaminergic neuron degeneration, which may result from autophagy impairment and α-Synuclein accumulation. First, we analysed the levels of TH and TUJ1 (Neuron-specific class III beta-tubulin), which is a neuronal marker, by Western Blot. Representative immunoblots are illustrated in Fig. 5a. Western blot analysis at 50 and 70 DoD showed no significant difference in TUJ1 levels between PD and control hMOs (Fig. 5b) but decreased levels of TH and lower TH to TUJ1 ratio in PD hMOs at 70 DoD (Fig. 5c, 5d), suggesting dopaminergic neurons loss at late stages of hMOs maturation. Then, we performed immunostaining for further analysis, with representative images of MAP2 and TH at 70 DoD are shown in Fig. 5e. Immunostaining confirmed fewer dopaminergic neurons in PD hMOs at 70 DoD (Fig. 5f) and increased fragmentation of TH-positive neurites in PD hMOs at 50 and 70 DoD (Fig. 5g).

**Figure 5.**
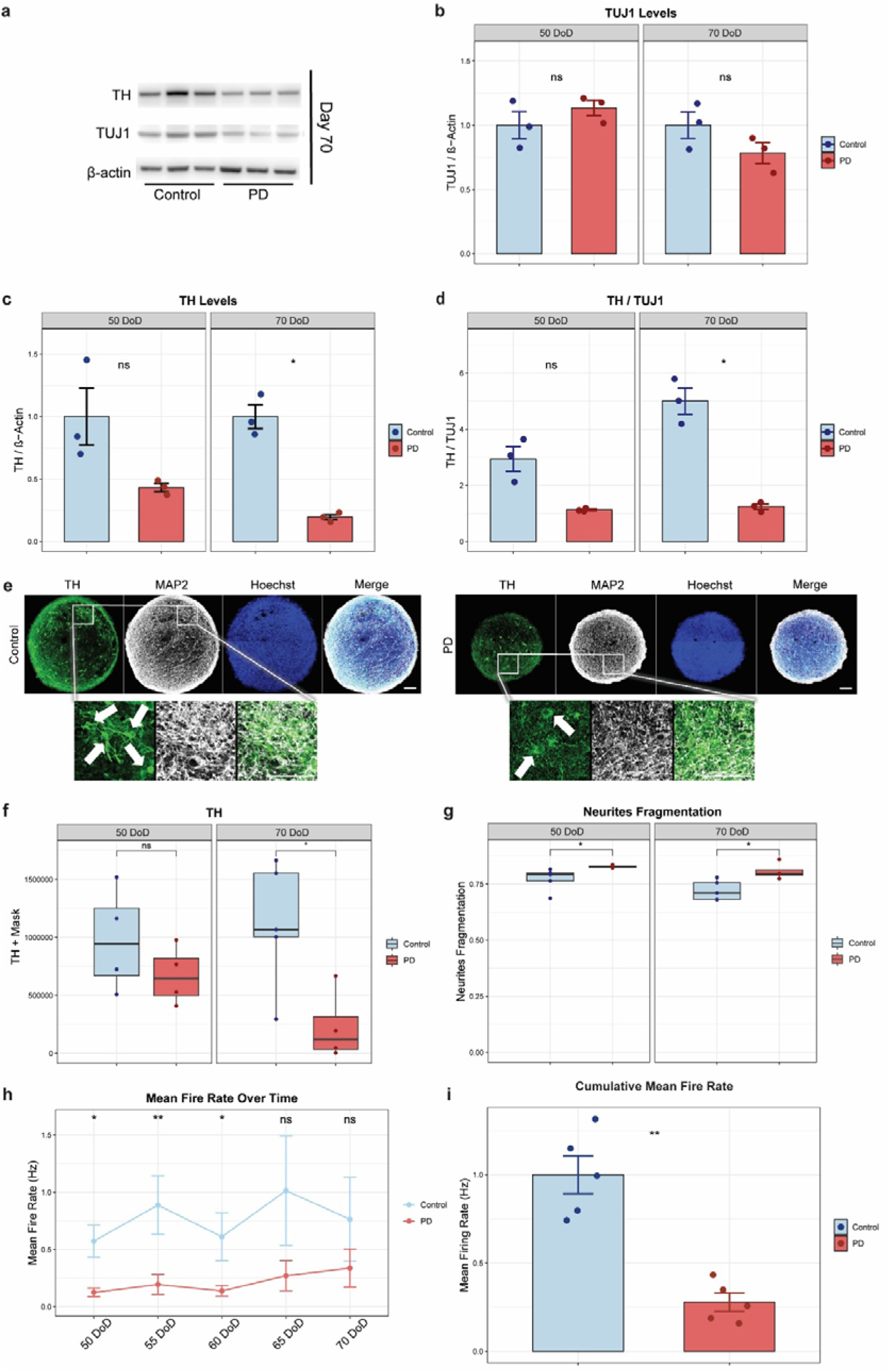
Progressive dopaminergic neuron degeneration and functional decline in PD hMOs. (a) Representative Western blot images showing TUJ1, TH, and β-Actin in control and PD hMOs. (b-d) Bar graphs depict protein levels of (b) TUJ1, (c) TH, and (d) the ratio of TH to TUJ1, showing a trend towards decreased dopaminergic neurons in PD hMOs. Protein expression was normalized to β-Actin. Immunoblot analysis was performed using ImageJ software. Error bars represent SEM. Data represent results from three independent experiments. Values are normalized to the average of controls at each time point. Statistical analysis was performed using Wilcoxon T-test. (e) Representative immunofluorescence images of hMOs stained for MAP2 (red), TH (green) and nuclei (blue). Scale bar: 20 μm. (f, g) Quantification of (f) TH-positive mask, (g) TH-positive neurites fragmentation, showing progressive dopaminergic neuron loss and morphological changes in PD hMOs. Data represent results from three independent experiments, each analyzed from at least two fields per hMO. Values are normalized to the average of controls across all time points, allowing a comparison over time. Statistical analysis was performed using Wilcoxon T-test. *p < 0.05, with “ns” indicating non-significant results. (h,i) (h) Mean fire rate of neurons in hMOs was measured every 5 days from 50 to 70 DoD. (i) Cumulative mean fire rate, showing decreased neuronal activity in PD hMOs. Data represent results from five independent experiments. Values are normalized to the average of controls at each time point. Statistical analysis was performed using Wilcoxon T-test; *p < 0.05, **p < 0.01, with “ns” indicating non-significant results. Error bars represent SEM. Image analysis was performed using MATLAB software for automated quantification of immunofluorescence staining.

Finally, we used Microelectrode array (MEA) to assess neuronal network functional activity in hMOs. Electrophysiological activity was recorded from 50 to 70 DoD, with intervals of 5 days. PD hMOs showed a decrease in the mean fire rate over time as well as in the cumulative mean fire rate compared to controls (Fig. 5h, i). Moreover, PD hMOs also tend to have less bursting events and lower cumulative burst frequency compared to control hMOs (see Supplementary Fig. 4). In summary, these results showed that in PD hMOs, dopaminergic neuron degeneration begins during the later stages of culture. Importantly, this degeneration seemed to affect dopaminergic neurons rather than other neurons since the levels of the general neuronal population remained unchanged. Furthermore, the reduced electrophysiological activity in PD hMOs precedes dopaminergic neuronal loss, suggesting that functional decline may be an early indicator of PD progression.

## Discussion

α-Synuclein aggregation and dopaminergic neuronal degeneration are the main neuropathological features of PD. Growing evidence suggest that autophagy dysregulation is a critical factor in PD pathology, particularly in the clearance of misfolded α-Synuclein. However, the timing and progression of autophagic processes are still poorly understood. This study explores the temporal dynamics of autophagy using live-cell imaging in two iPSCs-derived PD models, neuronal cultures and hMOs carrying the 3xSNCA mutation. Our results reveal that autophagy dysfunction emerges early in PD neuronal cultures, suggesting an early event in PD pathogenesis. In hMOs, while elevated α-Synuclein levels are already detectable at intermediate stages (50 DoD), full autophagy dysfunction becomes prominent only at later time points (70 DoD), supporting a context-dependent relationship between these pathological features.

Unlike traditional static analysis, the LC3-Rosella reporter system allows us to monitor autolysosomes in real time and therefore a better understanding of autophagy progression in PD, namely autophagosome maturation along with lysosome fusion. When integrating this strategy with biochemical analyses and with immunofluorescence analyses of α-Synuclein levels and dopaminergic markers, this strategy enabled a multidimensional perspective of disease progression.

Several studies have studied autophagy in iPSCs-derived dopaminergic neurons. In iPSCs-derived midbrain neurons carrying a 3xSNCA, LC3II and p62 flux was decreased and α-Synuclein impaired the autophagic flux shortly after autophagosome formation ^41^. Moreover, iPSCs-derived neurons from *GBA1*-PD and *LRRK2*-PD with the G2019S mutation also exhibited autophagy impairment with decreased autophagic flux and accumulation of autophagosomes ^42,43^. On the other hand, iPSCs-derived neurons carrying an *SNCA* triplication showed upregulated LC3II-autophagic flux, suggestive of induction of autophagy ^44^. A few studies have also investigated the role of autophagy in hMOs. Impairments in the autophagy flux of hMOs carrying a N370S mutation in the *GBA* gene were observed, where these hMOs showed decreased LC3II/LC3I ratio and increased levels of p62 ^45^. Also, DJ1 KO hMOs revealed a decrease in the LC3II flux showing a failure in the autophagy ^46^. Nevertheless, these studies were done in fixed samples, showing only a snapshot of a dynamic process. In our study, we assessed autophagy in live cells, providing a dynamic overview.

In PD neuronal cultures, autolysosome formation was reduced from the onset of differentiation. At 0 DoD, we observed a lower density of autolysosomes, especially of the small autolysosomes, compared to the control. Early in autophagy, an autolysosome grows as several lysosomes merge with an autophagosome ^38^. However, persistent accumulation of large autolysosomes is often associated with impaired degradation capacity and reduced autophagic flux. On the other hand, smaller autolysosomes are associated with more efficient and active degradation ^37,38^. As small autolysosomes are indicative of functional and active autophagic structures, these results suggest an early defect in autophagy initiation or autolysosome formation in PD neurons. At 3 DoD, we observed an increase in the autolysosome area in PD neurons accompanied by a decrease in the density of small autolysosomes but an increase in the density of large autolysosomes. Defects in the lysosome function also lead to an increase in the density of larger autolysosomes ^47^. While evaluating lysosomal activity may provide additional insights into these findings, it falls outside the scope of the present study and should be addressed in future investigations. As differentiation continued, PD neurons exhibited diminished autolysosome area and density regardless of their size, indicating a severe blockage of this pathway. This impairment strengthens the idea that, as PD develops, autophagy is not only compromised early but declines further.

Similar results were observed in PD hMOs showing a significant autophagy impairment. At 50 DoD, PD hMOs had a decrease in the autolysosome area. This came before the prominent impairment that was seen at 70 DoD, when autolysosome density and area were considerably reduced across all size groups, corroborating our observations in PD neuronal cultures at late stages of differentiation. Importantly, this autophagy dysfunction overlapped with increased accumulation of α-Synuclein, particularly in TH-positive neurons, in PD hMOs. Also, the increased ratio and colocalization of α-Synuclein with pS129 indicates the formation of aggregates. Mohamed *et al*. demonstrated that pS129 aggregates increased with the maturation of hMOs ^32^. Although these observations point toward an altered phosphorylation profile of α-Synuclein in PD hMOs, further investigation is needed to understand the exact nature of these pS129-positive structures.

To assess protein aggregation, we used additional markers such as p62, Proteostat, and Amytracker with α-Synuclein. Our results demonstrated elevated levels of p62 together with increased colocalization of α-Synuclein with Proteostat and Amytracker, and α-Synuclein with Proteostat and p62, confirming the presence of α-Synuclein aggregates resistant to degradation. It has been shown that α-Synuclein aggregates are resistant to degradation and inhibit autophagy by impairing autophagosome maturation ^48^. Also, α-Synuclein fibrils have been found to alter lysosome morphology, reducing the clearance of aggregates by autophagy ^49^. Although our results suggest a bidirectional link between autophagy and α-Synuclein levels, the autophagy system likely becomes overwhelmed with the increase of α-Synuclein levels.

We also explored how these impairments impact neuronal function. Neurons are particularly sensitive to changes in protein degradation pathways, with autophagy playing an important role in their survival ^50^. A few studies have demonstrated that α-Synuclein can lead to early pathological synaptic alterations in PD, where disruptions in dopaminergic neurotransmission have been documented to precede Lewy pathology and neurodegeneration ^51,52^. Thus, we assess the dopaminergic population and the electrophysiological activity of hMOs. In PD hMOs, a lower number of dopaminergic neurons at 70 DoD was observed. MEA analysis revealed that PD hMOs presented decreased neuronal activity. This functional decline correlated with autophagy dysfunction and α-Synuclein accumulation and aggregation, which altogether may be leading to dopaminergic degeneration. These results are corroborated by previous studies from our group showing that accumulation of α-Synuclein and decreased electrophysiological activity precede dopaminergic degeneration in PD hMOs carrying an 3xSNCA ^33,34^.

Although we report an interplay between α-Synuclein accumulation, autophagy dysfunction, and dopaminergic degeneration, additionally also other pathological processes are likely to be involved in PD progression. For instance, α-Synuclein oligomers triggered oxidative stress in iPSCs-derived neurons with a 3xSNCA ^53^. Also, α-Synuclein can induce mitochondrial dysfunction, calcium homeostasis disorders, endoplasmic reticulum stress and neuroinflammation ^54^. Exploring the role of these processes in autophagy in the context of 3xSNCA will provide a more thorough understanding of PD progression.

While our study provides valuable insights, it naturally comes also with limitations. The 3xSNCA mutation represents a specific genetic form of PD, potentially limiting translation our findings to other genetic backgrounds. Additionally, our here used hMO model, while more complex than neuronal cultures, still lacks critical cellular components, such as vascularization and immune components. For instance, the role of microglial autophagy has been described as a key factor in PD progression and pathogenesis ^55^. Choi *et al*. demonstrated that microglia have a neuroprotective effect by participating in the clearance of α-Synuclein ^56^. Moreover, longer studies in hMOs will provide a better understanding of late-stage PD pathology.

In conclusion, our study shows that autophagy dysfunction is a key characteristic of PD pathology. The integration of live-cell imaging and functional assays supports autophagy as a central player in PD pathogenesis and a promising target for early therapeutic intervention.

## Materials and Methods

### Ethical approval

This work with iPSCs has been approved by the Ethics Review Panel (ERP) of the University of Luxembourg and the national Luxembourgish Research Ethics Committee (CNER, Comité National d’Ethique de Recherche; CNER No. 201901/01).

### Generation and Maintenance of iPSCs, NESCs, Neuronal Cultures, and hMOs

#### iPSCs and NESCs Culture

Five iPSCs lines were used in this study, including two healthy controls and three PD lines carrying the 3xSNCA. A Rosella construct paired with the protein LC3 was integrated into one healthy control line and one PD line for autophagy analysis, as previously described ^35^ and in section 2.5. Detailed information about the cell lines can be found in Supplementary Table 1. iPSCs were cultured in Essential 8 medium (ThermoFisher Scientific, cat. no. A1517001) supplemented with 1% Penicillin/Streptomycin (P/S, Invitrogen, cat. no. 15140122) on Matrigel-coated plates (Corning, cat. no. 354277). Cells were passaged every 3-4 days using Accutase (Sigma-Aldrich, cat. no. A6964). After splitting, the medium was supplemented with 10 μM ROCK inhibitor Y-27632 (Merck Millipore, cat. no. 688000) for 24 h to enhance cell survival. All cultures were maintained at 37°C with 5% CO_2_.

NESCs were derived from iPSCs as previously described by Reinhardt et al., (2013) ^36^. The passage numbers of NESCs used for these experiments ranged from P10 to P14. NESCs were maintained in N2B27 medium on Matrigel-coated plates, supplemented with 0.75 μM purmorphamine (PMA; Enzo Life Sciences, cat. no. ALX-420-045), 3 μM CHIR-99021 (CHIR; Axon Medchem, cat. no. 1386), and 150 μM ascorbic acid (AA; Sigma-Aldrich, cat. no. A4403), referred to as N2B27 maintenance medium. N2B27 medium consisted of a 50:50 mixture of DMEM-F12 (Invitrogen; cat. no. 11320033) and Neurobasal (Invitrogen; cat. no. 21103049) with 1:200 N2 supplement (Invitrogen; cat. no. 17502048), 1:100 B27 supplement without Vitamin A (Invitrogen; cat. no. 12587010), 1% Glutamax (ThermoFisher Scientific, cat. no. 35050061), and 1% P/S. NESCs were passaged at 80–90% confluence using Accutase ^29^. Once the quality of the derivation was validated, neuronal cultures and hMOs were generated.

#### Neuronal Differentiation

For neuronal cultures, 3×10^4^ cells/well were seeded in Matrigel-coated cell carrier ultra plates (PerkinElmer, cat. no. 6055302). Neuronal differentiation was initiated by culturing NESCs in N2B27 maintenance medium for 2 days, followed by 6 days in N2B27 patterning medium supplemented with 200 μM AA, 500 μM Dibutyryl Cyclic Adenosine Monophosphate (db cAMP, Sigma-Aldrich, cat. no. D0627), 10 ng/mL human Glial cell line-derived Neurotrophic Factor (hGDNF, Peprotech, cat. no. 450-10), 10 ng/mL human Brain-Derived Neurotrophic Factor (hBDNF, Peprotech; cat. no. 450-02), 1 ng/mL Transforming growth factor beta 3 (TGF-β3; Peprotech; cat. no. 100-36E), and 1 μM PMA. The medium was changed every other day. Afterward, cells were cultured in N2B27 differentiation medium (identical to the patterning medium but without PMA) until the end of the experiments, totaling 11 DoD. Medium was changed every 3-4 days during this period. For the autophagy experiments, neuronal cultures were generated from NESCs containing the LC3-Rosella construct.

#### Organoid Generation

hMOs were generated as previously described ^29^, with slight modifications. A total of 9000 NESCs were seeded into each well of an ultra-low attachment 96-well round bottom plate (Corning, cat. no. 7007) and cultured in N2B27 maintenance medium for 2 days. For immunohistochemistry, western blot, and MEA analysis, hMOs were generated from regular NESCs. For the autophagy experiments, mosaic hMOs (55% LC3-Rosella NESCs + 45% regular) were generated to optimize signal resolution while reducing possible oversaturation, improving the autophagy measurements. The 3D colonies were then cultured in N2B27 patterning medium with medium changes every other day for 6 days. Differentiation was initiated with the N2B27 differentiation medium. To generate mosaic hMOs, the 3D colonies were embedded into 30 µl Geltrex droplets (Invitrogen, cat. no. A1413302), as previously described ^29^, placed into 24-well ultra-low adhesion plates (Corning, cat. no. 3473), and kept at 37°C and 5% CO_2_, under shaking conditions (80 rpm). For other experiments, 3D colonies were left in 96-well ultralow adhesion plates, at 37°C and 5% CO_2_, in static conditions. All hMOs were kept in culture for 50 and 70 DoD, with medium changes every 3-4 days.

### Immunofluorescence of NESCs and neuronal cultures

Cells were fixed with 4% paraformaldehyde (PFA, Electron Microscopy Sciences, cat. no. 15710) in phosphate-buffered saline (PBS) for 15 min at room temperature (RT), followed by three washes with PBS for 5 min each. Permeabilization was performed using 0.5% Triton X-100 in PBS for 15 min at RT, followed by blocking with 10% fetal calf serum (FCS) in PBS for 1 h at RT. Cells were then incubated overnight at 4°C with primary antibodies (Supplementary Table 2) diluted in 3% FCS in PBS. After three 5-min washes with PBS, cells were incubated with secondary antibodies (Supplementary Table 2) and Hoechst 33342 (1:10000, Life Technologies, cat. no. H3570) for nuclear staining, diluted in 3% FCS in PBS, for 1 h at RT in the dark. Three additional 5-min washes with PBS were performed.

For NESC characterization, cells were mounted with Fluoromount-G mounting (Southern Biotech, cat. no. 0100-01) and imaged using a Zeiss LSM 710 Confocal Inverted Microscope (*RRID:SCR_018063*) with a total magnification of 200x.

For other stainings, cell carrier ultra plates with duplicates per condition from 3 independent experiments were imaged in an automated manner using the Yokogawa CellVoyager CV8000 microscope (*RRID:SCR_023270*) with a 20x objective.

### Immunofluorescence of hMOs

hMOs were fixed overnight with 4% PFA in PBS at 4°C and washed three times with PBS for 15 min each at RT. One hMO per condition and independent experiment was embedded in 3% low-melting point agarose (Sigma-Aldrich, cat. no. A9414) in PBS. The agarose block was sectioned using a vibrating blade microtome (Leica VT1000s, *RRID:SCR_016495*) into 50 µm sections that were collected sequentially into three wells of a 48-well plate. Sections were permeabilized and blocked for 2 h at RT in blocking buffer (2.5% normal goat or donkey serum, 2.5% bovine serum albumin, 0.1% sodium azide, and 0.5% Triton X-100 in PBS) with shaking. Sections were then incubated with primary antibodies (Supplementary Table 2) diluted in blocking buffer containing 0.1% Triton X-100, for 48 h at 4°C with gentle shaking. Then, sections were washed 3x with 0.01% Triton X-100 in PBS for 10 min each and subsequently incubated with the secondary antibodies (Supplementary Table 2), along with Hoechst 33342 (1:10000 dilution), diluted in blocking buffer containing 0.1% Triton X-100, for 2 h at RT with shaking. Sections were washed three times with 0.01% Triton X-100 in PBS for 10 min each and once with Milli-Q water. For protein aggregates, we used the Proteostat® Aggresome Detection Kit (Enzo, cat no. ENZ-51035) and Amytracker 680 (Ebba Biotech AB) protocols. Any remaining agarose was carefully removed before mounting the sections with Fluoromount-G mounting medium. Sequential collection and staining of sections were implemented to avoid bias in selecting specific hMO regions.

Imaging of the sections from at least 3 independent hMOs per condition was performed using a Yokogawa CellVoyager CV8000 high-content screening microscope. For α-Synuclein aggregates detection, z-planes were acquired using a 60x oil objective. For other high-content image analyses, sections were imaged with a 20x objective.

### LC3-Rosella Autophagy Reporter

The LC3-Rosella construct, which combines the pH-sensitive green fluorescent protein pHluorin and the red fluorescent protein from Discosoma (DsRed) with LC3II, was integrated into iPSCs lines via nucleofection, using the P3 Primary Cell 4D-Nucleofector® X Kit L (Lonza Bioscience, cat. no. V4XP-3024), as previously described ^35,57^. Transfected iPSCs were sorted by fluorescence-activated cell sorting (FACS) to enrich positive clones and screened by fluorescence microscopy to confirm LC3-Rosella expression. The use of pH-sensitive fluorescent proteins enables the distinction of several subcellular compartments. This happens because the pHluorin signal, a modified form of GFP, is suppressed in acidic conditions, such as the interior of the autolysosome, while DsRed continues to fluorescence. However, since LC3 is a main component of the neuronal cytoskeleton ^58^, pHluorin signal is very diffuse in neuronal cultures and hMOs. Hence, our analysis focused on autolysosomes identified as red-only vesicles (pHIuorin-DsRed+ structures). For imaging, pHluorin and DsRed were excited using 488 nm and 561 nm lasers, respectively.

### Live Imaging in Neuronal Cultures

Autophagy dynamics were monitored in live neuronal cultures at 0, 3, 5, 7, 9, and 11 DoD using the Yokogawa CV8000 microscope, under controlled atmosphere conditions (37°C and 5% CO_2_). Cell carrier ultra plates with triplicates per condition from 3 independent experiments were imaged, acquiring a total of 25 fields per well. DsRed and pHluorin images were acquired in parallel using camera binning 2 and a 20x objective. Cells were kept under normal conditions between imaging sessions.

### Live Imaging of hMOs

Live imaging of LC3-Rosella in hMOs was performed at 50 and 70 DoD using the Zeiss Cell Observer Spinning Disk confocal microscope, under controlled atmosphere conditions (37°C and 5% CO_2_). Individual hMOs from 3 independent experiments were transferred to a µ-Slide 8 Well Chambered Coverslip (Ibidi, cat. no. 80826) and allowed to attach for 45 min without medium in the incubator. Then, N2B27 maturation medium was added, and hMOs were imaged live. DsRed and pHluorin signals were imaged in parallel using a 40x oil objective. A total of 3 fields per hMO were acquired. After imaging, hMOs were then returned to culture for further analysis.

### Image processing and Autolysosome Classification

Images of neuronal cultures and hMOs were processed and analyzed using custom algorithms developed in MATLAB (2020a, Mathworks; *RRID:SCR_001622*), as described by our group ^29,35^. These in-house imaging analysis pipelines allow an automated identification and segmentation of cellular structures, such as nuclei, neurons and autophagosomes by extracting their specific features.

For the autophagy experiments, autolysosome size was classified according to voxel area, based on histogram analysis of their distribution in each model. In neuronal cultures, autolysosomes were defined as small (<8 voxels, corresponding to <0.09 µm²), medium (8 to 25 voxels, corresponding to 0.09–0.29 µm²), and large (≥25 voxels, corresponding to ≥0.29 µm²). In mosaic hMOs, given their greater structural complexity and signal distribution, we adjusted the thresholds to small (<25 voxels, <0.29 µm²), medium (25 to 80 voxels, 0.29–0.93 µm²), and large (≥80 voxels, ≥0.93 µm²). These µm² values were calculated based on the voxel dimensions from the imaging system. This size-based classification allowed us to do a more detailed analysis of autophagy.

### Western Blot

Five hMOs per condition were lysed using RIPA buffer (Abcam, cat no. ab156034) supplemented with protease inhibitor cocktail (Roche, cat no. 11697498001) and phosphatase inhibitor cocktail (Merck Millipore, cat no. 524629). To improve protein extraction and sample quality, lysates were sonicated for 10 cycles (30 s on/30 s off) using the Bioruptor Pico (Diagenode) and centrifuged at 14,000 x g for 20 min at 4°C. Protein concentrations were determined using the Pierce™ BCA Protein Assay Kit (ThermoFisher Scientific, cat no. 23225). Protein samples (10 µg) were prepared and boiled at 95 °C for 5 min in loading buffer. Proteins were separated on 4-12% NuPAGE™ Bis-Tris polyacrylamide gels (ThermoFisher Scientific), using MES SDS running buffer (ThermoFisher Scientific), at 110 V for 1 h 40 min. Proteins were transferred to polyvinylidene difluoride (PVDF) membranes using the iBlot™ 2 Gel Transfer Device (ThermoFisher Scientific), following the manufacturer’s instructions. Membranes were fixed in 0.04% PFA for 30 min at RT and washed three times with 0.01% Tween in PBS (PBS-T) for 5 min each. Then, membranes were blocked in 5% non-fat powdered milk (Roth, cat. no. T145.2) prepared in 0.02% PBS-T for 1 h at RT, followed by overnight incubation at 4°C with primary antibodies (Supplementary Table 2) diluted in 5% BSA in 0.02% PBS-T. After three 5-min washes with 0.01% PBS-T, membranes were incubated with HRP-linked secondary antibodies (anti-rabbit IgG (H + L) or anti-mouse IgG (H + L), VWR) at a 1:1000 dilution in 0.02% PBS-T for 2 h at RT. Membranes were then washed three times with 0.01% PBS-T 5 min each, and protein bands were detected using SuperSignal West Pico Chemiluminescent Substrate (ThermoFisher Scientific, cat. no. 34580) and imaged with the STELLA 8300 imaging system (Raytest). Image analysis was performed using ImageJ software (NIH; *RRID:SCR_003070*). Band intensities were normalized to β-actin as a loading control. Samples from the same independent experiment were loaded onto a single gel, and all blots were processed using the same experimental conditions. Western blot experiments were conducted from 3 independent experiments per condition.

### MEA Analysis

MEA recordings were performed using the Maestro system (Axion BioSystems) to record the spontaneous electrical activity of hMOs, from 50 to 70 DoD, with intervals of 5 days. Before the recordings, 48-well MEA plates (16 electrodes per well) were coated with 0.1 mg/mL poly-D-lysine hydrobromide (Sigma, cat. no. P7886) and 10 μg/mL laminin (Sigma, cat. no. L2020) at 37°C. Then, 45-day-old hMOs were carefully positioned onto the electrode arrays and secured with a droplet of Geltrex (Invitrogen, cat. no. A1413302). Following a short polymerization period, N2B27 differentiation medium was added. 24 h before each recording session, fresh N2B27 differentiation medium was added to the hMOs.

MEA recordings were performed as previously described ^29,34,59^. Spontaneous activity was recorded for 10 minutes at 37°C. Data acquisition and analysis were performed using Axion Integrated Studio (AxIS 2.1; *RRID:SCR_016308*) software. To minimize false positives and missed detections, a Butterworth band-pass filter (200-3000 Hz) and a threshold of 6x standard deviation were applied. Electrodes registering an average of ≥5 spikes/min were considered active. Neural stat compiler files were used for data analysis in MATLAB. The parameters analyzed were spike frequency and burst analysis. This non-invasive approach allowed for longitudinal monitoring of hMOs maturation and network development over time. For each condition, hMOs were analyzed in each of the 5 independent experiments.

### Statistical analysis

Data process and statistical analyses were performed using R software (R version 4.3.0 - “Already Tomorrow”, *RRID:SCR_001905*). For each cell line and experiment, data were grouped by condition, with mean values calculated. All experiments were performed with a minimum of 3 biological replicates. The normality test was done by using the Shapiro-Wilk test. Statistical comparisons between control and experimental groups (Control vs. PD) were conducted using the Wilcoxon signed-rank test or t-test, depending on data distribution. Significance levels are denoted as p < 0.05 (*), p < 0.01 (**), p < 0.001 (***), and p < 0.0001 (****), with “ns” indicating non-significant results. Plots were generated using R Studio, and sample sizes, replicates, and experimental batches are detailed in figure legends.

## Data availability

All data generated or analysed during this study as well as scripts that support the findings are available at 10.17881/gd2k-9685.

## Code availability

All scripts used to obtain, analyze and plot the data are available at https://gitlab.com/uniluxembourg/lcsb/developmental-and-cellular-biology/almeida_2025.

## Acknowledgements

The authors acknowledge the private donors supporting research at the Luxembourg Centre for Systems Biomedicine (LCSB). Microscopy work was supported by the LCSB Bioimaging Platform. We thank Dr. Nathasia Mudiwa Muwanigwa and Dr. Jennifer Modamio Chamarro for their contributions to the characterization of iPSCs and derivation of NESCs used in this study.

This work was funded by FCT – Fundação para a Ciência e a Tecnologia, I.P. (Portugal) through doctoral grant SFRH/BD/149036/2019 (DOI: 10.54499/SFRH/BD/149036/2019) and COMPETE2030-FEDER-00795500 (n°16772).

The use of existing iPSCs lines obtained from previous studies was approved by the Luxembourg National Ethics Committee (Comité National d’Éthique de Recherche, CNER No. 201901/01). Informed consent was obtained from all individuals prior to sample donation, in accordance with approved protocols and written consent forms.

## Author contributions

CS-A conceived and designed the study, collected data, performed data analysis and interpretation. JJ assisted with automated image analysis. GG-G, IR, AZ and EZ collected part of the stainings data. GG-G, IR, CS and DFr contributed to the experimental work. DF assisted with MEA analysis. The study was supervised by JCS. The original manuscript was written by CS-A and ACC, LB and JCS reviewed and edited the manuscript. All authors read and approved the final manuscript.

