## Supplementary Data for "Autophagy Dysfunction in iPSCs-Derived Neurons and Midbrain Organoids Carrying a *SNCA* Triplication"

***Supplementary Figure 1***

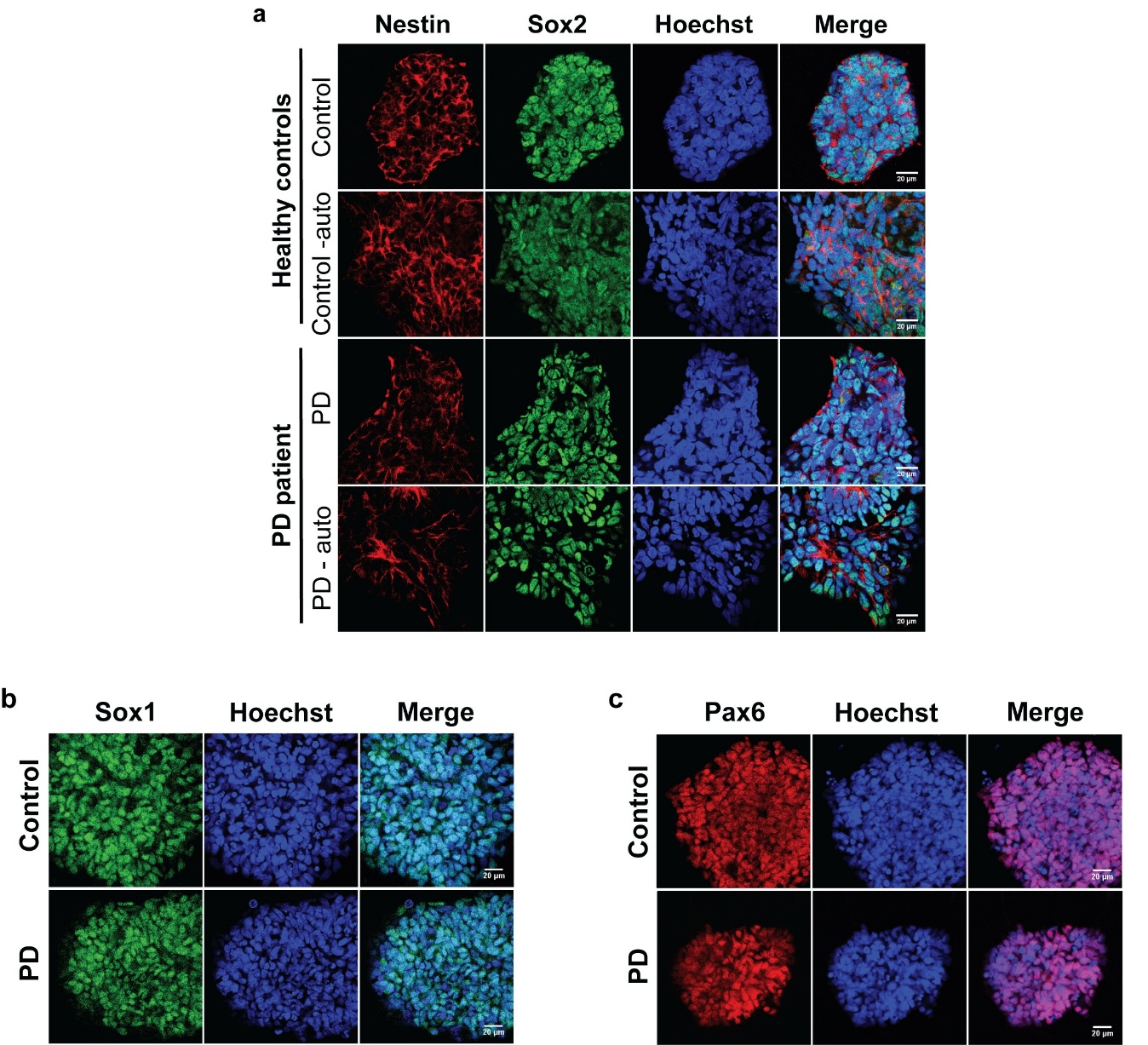

**Supplementary Figure 1 - Neuroepithelial stem cells (NESCs) characterization.** a) Representative immunostainings of Control, Control – auto, PD and PD – auto NESCs, showing expression of Nestin and Sox2. b,c) Representative immunostainings of Control and PD NESCs, showing expression of b) Sox1 and c) Pax6. Scale bar: 20 µm.

***Supplementary Figure 2***

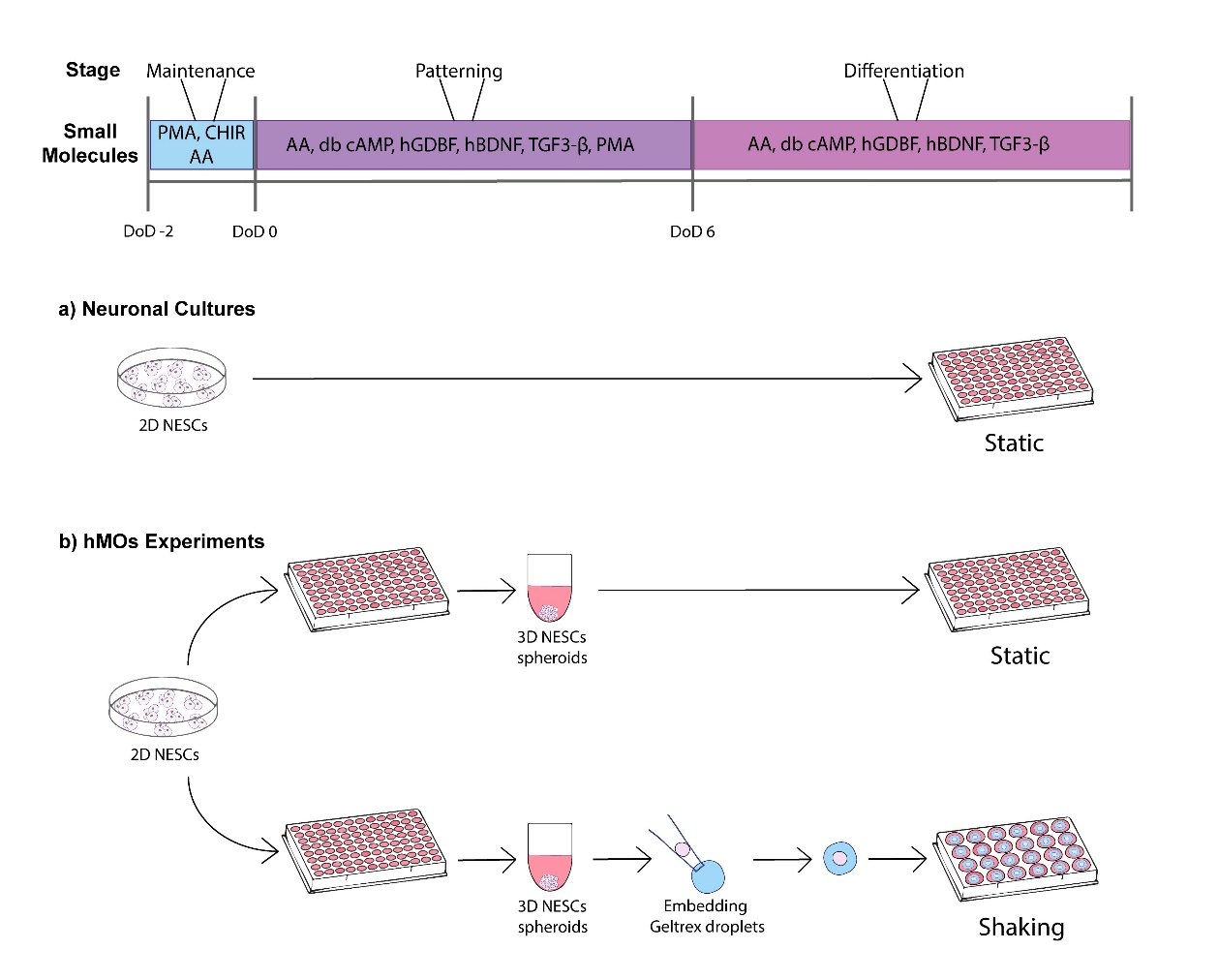

**Supplementary Figure 2 - Schematic overview of the protocol used for the generation of neuronal cultures and hMOs.**

***Supplementary Figure 3***

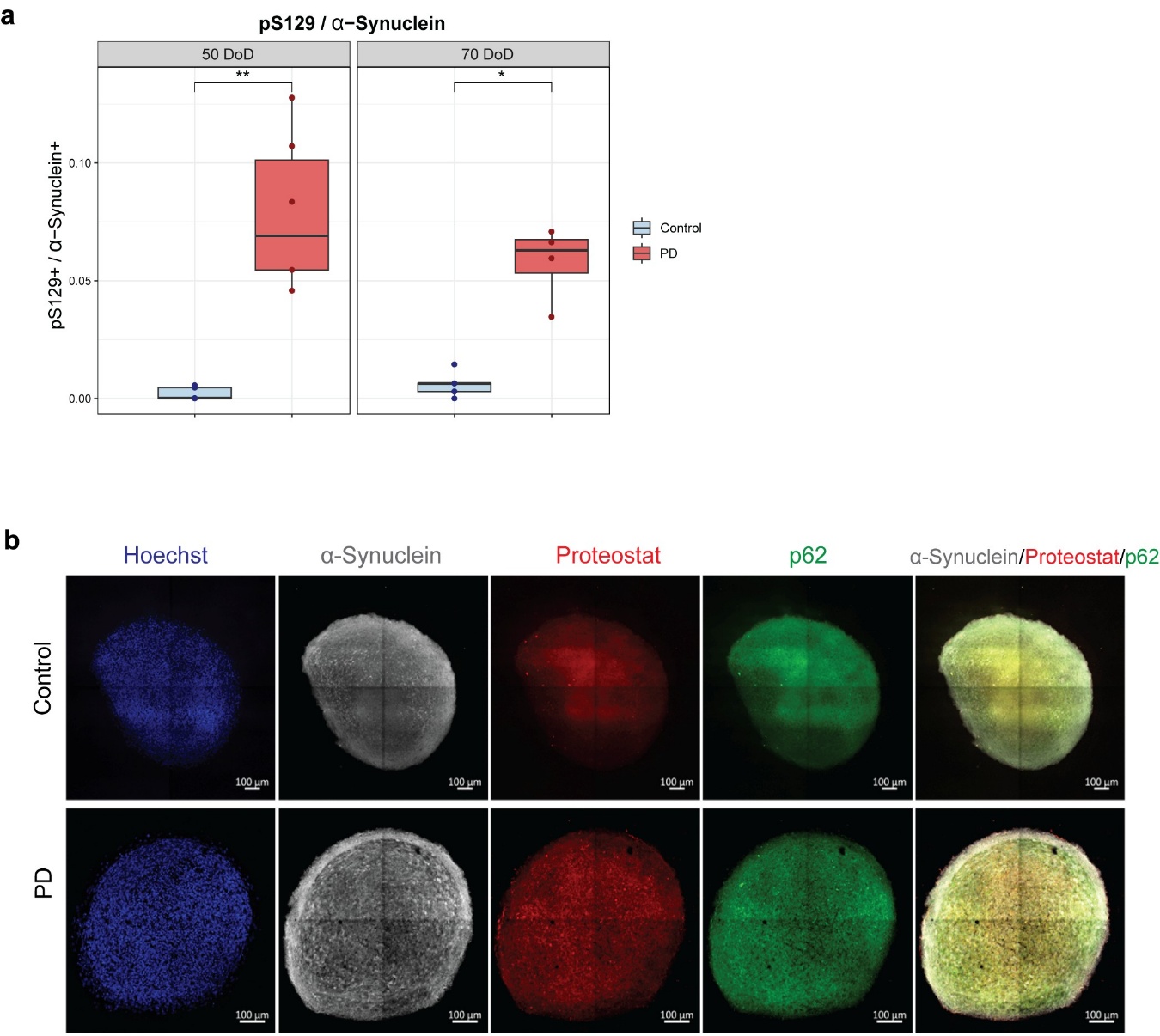

**Supplementary Figure 3 - PD hMOs exhibit elevated pS129 levels and increased α-Synuclein aggregation.** a) Ratio of pS129 / α-Synuclein in PD and control hMOs at 50 and 70 days of differentiation (DoD), analyzed by WB. b) Representative immunofluorescence images of 70 DoD hMOs stained for nuclei (blue), α-Synuclein (grey), Proteostat (red) and p62 (green). Scale bar: 100 µm.

***Supplementary Figure 4***

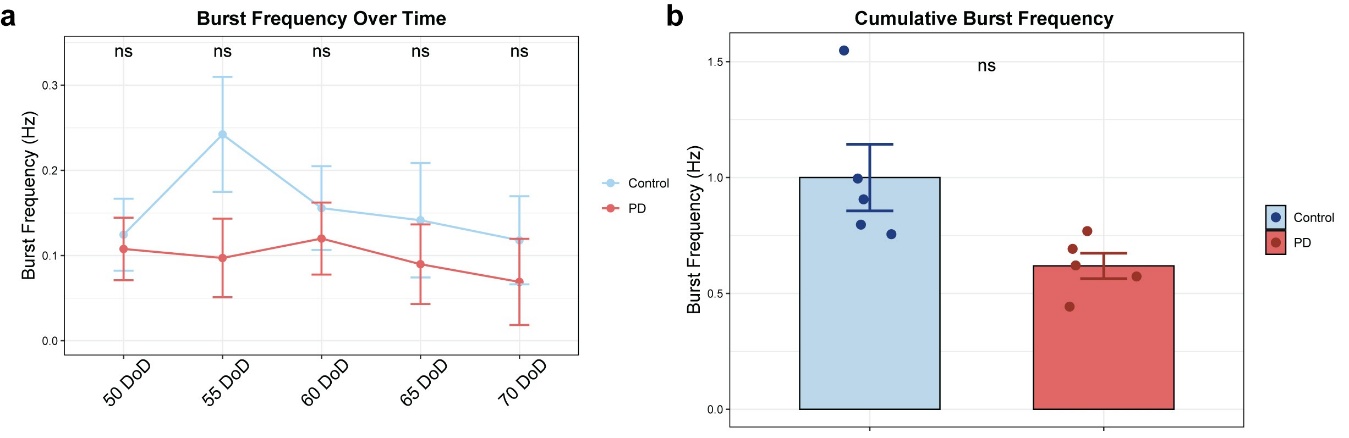

**Supplementary Figure 4 - PD hMOs exhibit a non-significant trend toward reduced burst dynamics.** a) Burst frequency in PD and control hMOs recorded every 5 days between 50 and 70 DoD. b) Cumulative burst frequency over the same period, suggesting lower overall neuronal activity in PD hMOs. Data represent the average of five independent experiments. Values were normalized to the mean of controls at each time point. Statistical analysis was performed using the Wilcoxon test, with "ns" indicating non-significant results. Error bars represent SEM. Burst detection and analysis were performed using automated MATLAB-based tools.

Supplementary Table 1. Human iPSCs used in this study

| **Cell Line Name** | **Autophagy sensor LC3-Rosella** | **Healthy (WT) or Patient (PD) Origin** | **Sex** | **Age of sampling** | **Age of onset** | **Source** | **Ref #** | **Karyotype** |
| --- | --- | --- | --- | --- | --- | --- | --- | --- |
| Control | No | WT | F | 55 | - | Reinhardt et al. 2013 |  | Normal |
| PD | No | PD | F | 55 | 50 | EBISC - EDi001-A | EDi001-A | Normal |
| SNCA KO | No | PD | F | 55 | 50 | Chen et al. 2019 | AST23-4KO-5B | Normal |
| Control - auto | Yes | WT | F | Cord  Blood  derived | - | GIBCO | A13777 | Normal |
| PD - auto | Yes | PD | F | 55 | 50 | EBISC - EDi001-A | EDi001-A | Normal |

Supplementary Table 2. Primary antibodies used in this study

| **Antibodies** | **Species** | **Source** | **Reference** | **RRID** | **WB Dilution** | **IF Dilution** |
| --- | --- | --- | --- | --- | --- | --- |
| α-Synuclein | Rabbit | Santa Cruz | sc7011-R | AB_2192953 | 1:1000 | 1:1000 |
| α-Synuclein (2A7) | Mouse | NOVUS  Biologicals | NBP1-05194 | AB_1555287 | - | 1:1000 |
| β-Actin | Mouse | Cell Signaling | 3700 | AB_2242334 | 1:50.000 | - |
| MAP2 | Chicken | Abcam | ab92434 | AB_2138147 |  | 1:1.000 |
| Nestin | Mouse | Millipore | MAB5326 | AB_2251134 | - | 1:100 |
| Pax6 | Rabbit | Covance | PRB-278P | AB_291612 | - | 1:300 |
| Phospho-α-Synuclein  (Ser129)  (D1R1R) | Rabbit | Cell Signaling | 23706S | AB_2798868 | 1:500 | 1:500 |
| p62  (SQSTM1) | Mouse | Abcam | ab56416 | AB_945626 | - | 1:500 |
| Sox1 | Goat | R&D Systems | AF3369 | AB_2239879 | - | 1:200 |
| Sox2 | Goat | R&D Systems | AF2018 | AB_355110 | - | 1:200 |
| TH | Rabbit | Abcam | ab112 | AB_297840 | 1:1.000 | 1:1.000 |
| TUJ1 | Mouse | BioLegend | 801201 | AB_2313773 | 1: 50.000 |  |
